# A generative model of the connectome with dynamic axon growth

**DOI:** 10.1101/2024.02.23.581824

**Authors:** Yuanzhe Liu, Caio Seguin, Richard F. Betzel, Danyal Akarca, Maria A. Di Biase, Andrew Zalesky

## Abstract

Connectome generative models, otherwise known as generative network models, provide insight into the wiring principles underpinning brain network organization. While these models can approximate numerous statistical properties of empirical networks, they typically fail to explicitly characterize an important contributor to brain organization – axonal growth. Emulating the chemoaffinity guided axonal growth, we provide a novel generative model in which axons dynamically steer the direction of propagation based on distance-dependent chemoattractive forces acting on their growth cones. This simple dynamic growth mechanism, despite being solely geometry-dependent, is shown to generate axonal fiber bundles with brain-like geometry and features of complex network architecture consistent with the human brain, including lognormally distributed connectivity weights, scale-free nodal degrees, small-worldness, and modularity. We demonstrate that our model parameters can be fitted to individual connectomes, enabling connectome dimensionality reduction and comparison of parameters between groups. Our work offers an opportunity to bridge studies of axon guidance and connectome development, providing new avenues for understanding neural development from a computational perspective.

**Author Summary:** Generative models of the human connectome provide insight into principles driving brain network development. However, current models do not capture axonal outgrowth, which is crucial to the formation of neural circuits. We develop a novel generative connectome model featuring dynamic axonal outgrowth, revealing the contribution of microscopic axonal guidance to the network topology and axonal geometry of macroscopic connectomes. Simple axonal outgrowth rules representing continuous chemoaffinity gradients are shown to generate complex, brain-like topologies and realistic axonal fascicle architectures. Our model is sufficiently sensitive to capture subtle interindividual differences in axonal outgrowth between healthy adults. Our results are significant because they reveal core principles that may give rise to both complex brain networks and brain-like axonal bundles, unifying neurogenesis across scales.

## Introduction

The network of axonal connections comprising a nervous system is known as the connectome (Hagmann, 2005; Sporns et al., 2005). Connectomes display non-random topological characteristics, such as small-worldness and modularity (Bassett & Bullmore, 2017; Sporns & Betzel, 2016) as well as rich diversity in the strength of connections and regions (Buzsáki & Mizuseki, 2014). While connectome topological properties are well characterized, the underlying wiring principles that give rise to these properties are poorly understood.

Generative models offer one avenue to investigate principles governing connectome development. Connectome-like networks have been generated *in silico* to model the micro-, meso-, and macro-scale neural connectivity of many organisms, including *C. Elegans*, *Drosophila*, non-human mammals, and humans, through a variety of spatial, topological, and physiological wiring rules (Akarca et al., 2023; Betzel et al., 2016; Beul et al., 2018; Ercsey-Ravasz et al., 2013; Faskowitz et al., 2018; Henriksen et al., 2016; Kaiser & Hilgetag, 2004; Klimm et al., 2014; Oldham et al., 2022; Pavlovic et al., 2014; Priebe et al., 2017; Simpson et al., 2011; Vértes et al., 2012). In each of these models, the extent to which the generative process is guided by each wiring rule is determined by a set of tunable parameters. Typically, connections are more likely to be generated between regions that are close in spatial proximity to each other (Ercsey-Ravasz et al., 2013; Kaiser & Hilgetag, 2004) and/or for which inclusion of the proposed connection would enhance a desired topological criterion (Simpson et al., 2011; Vértes et al., 2012).

Axonal growth and guidance are important mechanisms that shape brain wiring. Since Ramon y Cajal’s discovery of growth cones and Sperry’s pioneering chemoaffinity hypothesis (Cajal, 1890; Chilton, 2006; Sperry, 1963; Zang et al., 2021), a variety of guidance molecules, such as netrins and slits, were found to contribute to axon pathfinding (Brose et al., 1999; Kennedy et al., 1994; Kidd et al., 1999; Serafini et al., 1994). Miswired connectomes are evident in model organisms deficient in guidance molecules and receptors; for example, abnormal optic chiasm development has been found in slit-deficient mice (Dickson, 2002). Given the importance of axonal guidance in brain network development and wiring, modeling of pathfinding mechanisms may lead to improved connectome generative models that reflect multiple spatial phenomena, compared preferential generation of connections between pairs of regions in close spatial proximity. Connectome generative models that consider axonal guidance may provide insight into connectome development, complementing the insight provided by current models and shedding light on the mechanisms that generate the characteristic geometry and spatial architecture of axonal fiber bundles.

Explicitly simulating axonal growth also provides an opportunity to generate weighted brain networks. Most established connectome generative models are unweighted – a connection is either present or absent between a pair of regions. As such, information about diverse connectivity strengths is overlooked and not modeled. Recent studies have proposed various methods to address this issue, such as through connectome community (Faskowitz et al., 2018) and communicability redundancy (Akarca et al., 2023). Despite the unique strengths of these approaches, axon counts remain a natural and straightforward representation for the strengths of physical neural connectivity. In this direction, researchers have established connectome generative models that simulate networks by growing axons in predetermined directions (Song et al., 2014). In contrast to tuning weights of connections themselves, this work parcellates a continuous space into discrete regions that are connected by multiple simulated axons. As a result, connection weights naturally arise from the axon counts between pairs of regions. Although a nodal correspondence between generated and empirical connectomes is missing, this approach has the advantage of being biologically tractable. The generated networks are found to replicate many topological properties of empirical connectomes, including degree, clustering, and triad distributions (Song et al., 2014). However, it remains unclear whether these attributes persist or whether new topological characteristics arise in the presence of dynamic axon guidance.

In this study, we establish a new spatially embedded generative model for weighted connectomes. Our model significantly builds on the seminal models of Kaiser and colleagues (2009) as well as Song and colleagues (2014), both of which feature axon outgrowth. *Dynamic* axon growth is a key novelty of our model, without which curved axons cannot form. Each brain region exerts a distance-dependent attractive force on an extending axon’s tip, steering the direction of axon growth. This emulates the process through which axon growth cones react to molecular guiding cues, whose concentration decays with the distance to chemical release sites. We find that our model can recapitulate a diverse array of topological features characteristic of nervous systems, at the edge, node, and network levels. We fit the two parameters of our generative model to individual connectomes, generating weighted networks that reflect interindividual variations in brain network architecture. Overall, our work enables generation of connectomes *in silico* that are weighted, spatially embedded, and feature axonal trajectories that appear biologically realistic.

## Results

We develop a model that generates weighted connectomes through dynamic axon outgrowth, an extension of the static generative model proposed by Song et al. (2014). Specifically, the direction of axon growth in a static model is governed in a one-shot manner where fibers are generated in a direct, linear trajectory towards their targets. In contrast, axons in a dynamic model continuously assess the guidance gradient acting on them and respond accordingly. As such, dynamic models have an internal quasi-temporal structure such that the position of the axon in the past influences its subsequent position over iterations. As a result, axons are generated with richer, fascicle-like geometry.

For model simplicity and axon visualization purpose, we simplify the cerebral volume to a two-dimensional circular construct of radius *R*. The circumference and the internal space of the circle represent brain gray and white matter, respectively (Fig. 1a). To discretize the circle into regions, we uniformly divide the circumference into *N*_*n*_ segments of equal length, where each segment represents a distinct brain region, otherwise known as a *node*. Given that brain regions differ in volume and surface area, geometric heterogeneity of nodes is introduced through a parameter *ρ* that perturbs node center coordinates (Fig. 1b, detailed in Methods).

**Figure 1.**
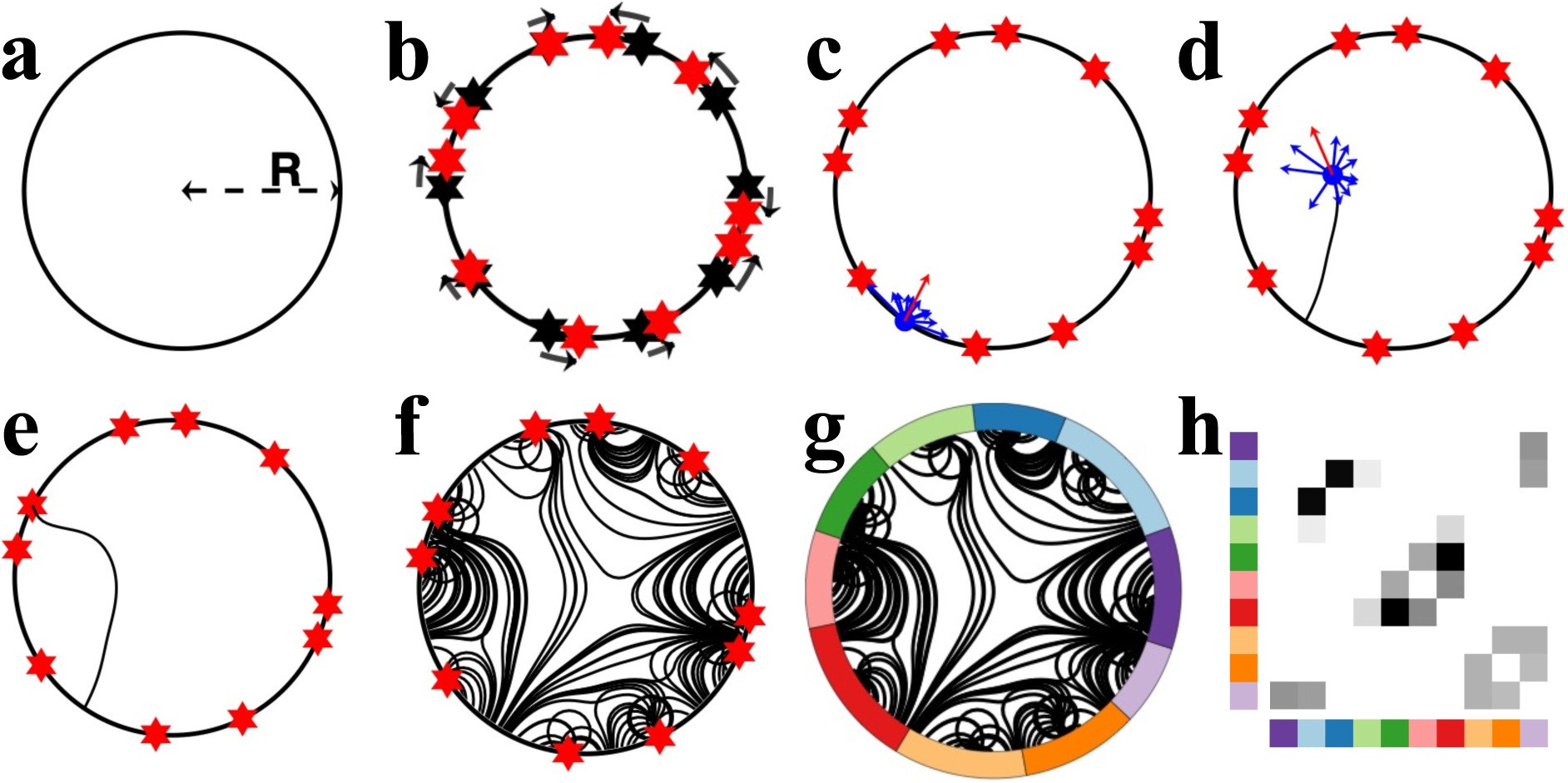
Illustrative example of the generative process governing axonal growth and network formation. **a)** The model is formulated on a circle of radius R. **b)** Coordinates are uniformly positioned along the circle’s circumference, each representing a node center (black hexagrams). We used ten nodes in this illustrative example. The coordinates are randomly perturbed on the circumference (in the direction of black arrows; new node centers are represented with red hexagrams) to introduce nodal heterogeneity. **c)** An axon is seeded on the perimeter. It perceives an attractive force from all nodes (blue arrows) and propagates step-by-step in the direction of the net force (red arrow). **d)** The net force experienced by the growth cone is updated at each propagation step to ensure nodes that become closer to the growth cone exert greater force, while nodes further from the growth cone exert less force. **e)** The simulated axon forms a connection when its growth cone reaches a point on the circular circumference. **f)** Multiple axons are generated, giving rise to structures resembling axonal fiber bundles. **g)** The endpoints of axons are assigned to the nearest nodes to construct a network. **h)** The generated network is represented using a weighted, undirected connectivity matrix.

We use the term *axon* to denote a unitary connection between a pair of nodes and use the term *growth cone* to refer to an axon’s growing tip. In our model, *N*_*a*_ axons are uniformly seeded at random from the circular circumference. Each axon is then propagated step-by-step within the circle’s interior until reaching a point on the circumference. Crucially, at each propagation step, the axon’s direction of propagation is updated based on a combined attractive force exerted by each node (Fig. 1c). The attractive forces can represent various environmental and molecular cues (Wadsworth, 2015), decaying as a function of distance between the node exerting the force and an axon’s growth cone (Kaiser et al., 2009; Murray, 2002). Specifically, we assume that the net force exerted at position 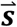 is given by:

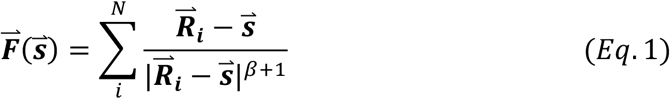

where 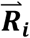 is the coordinate of the *it*ℎ node center, and |*x*| describes the vector magnitude of *x*. The parameter *β* regulates the power-law decay of attractive forces based on the distance between the growth cones and node centers. A larger *β* penalizes the attractive forces from distant nodes and promotes the formation of local connectivity. It is one of the two tunable parameters of the model.

A distinguishing feature of the model is dynamic axonal growth, echoing *in vitro* and *in vivo* evidence suggesting that axons actively modify growth pathways in response to local molecular and mechanical cues (Dickson, 2002; Oliveri & Goriely, 2022). The simulated axons grow progressively in a step-by-step manner, based on the attractive forces described by 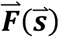 (Fig. 1d). For each step, an axon growth cone situated at position 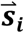 is extended in the direction of 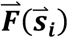 for a constant distance *L*_*s*_. This step length *L*_*s*_ is the second tunable parameter of the model. We additionally impose weak regularity constraints on growth (see Methods and Fig. S1) to avoid trajectories with biologically unrealistic sharp turns (Katz, 1985).

Axon propagation terminates as soon as its growth cone intersects with the circle, forming a connection between two points on the circle’s circumference (Fig. 1e, 1f). Axons that successfully connect two points of the circumference are selected for network construction. Some axons fail to navigate to a point on the circumference, and they are excluded from subsequent analyses (Methods and Fig. S2).

We construct networks by assigning the endpoints of successful axons to their nearest node centers (Fig. 1g, 1h). The connectivity weight between a pair of nodes is given by the total number of axons linking them. Despite the intrinsic directionality of simulated axons, for simplicity, we focussed on mapping weighted, undirected networks. Within-region connections are ignored.

### Generating brain-like axonal fiber bundles, hubs, and connectivity weights

In this study, unless otherwise specified, we matched *N*_*n*_ to the number of brain regions in the Desikan-Killiany atlas (84 nodes). It should be noted that nodes in our generated and empirical connectomes do not have a one-to-one correspondence, an intrinsic limitation from using simplified brain geometry (Song et al., 2014). We started by investigating how simulated networks behaved in response to variations in model parameters. Our model is governed by two key parameters (explained in Supplementary materials – Parameter specification and Fig. S3): *β* - the distance-dependent decay of the attractive force, and *L*_*s*_ - the length of each extending step. Note that *β* determines the relative contribution of guiding cues exerted by each node, such that a larger *β* emphasizes the guidance from local, adjacent nodes, relative to distant nodes; whereas *L*_*s*_ governs the extent to which an axon can change its trajectory per unit length.

We first evaluated the generated networks in terms of the connectivity weight and nodal degree distributions. As spatially embedded networks, connectomes exhibit strong and abundant short-range connections and weak and rare long-range connections across a variety of scales and species. This property is typically modeled using an exponential distance rule (EDR) of connection weights (Betzel & Bassett, 2018; Ercsey-Ravasz et al., 2013; Gămănuţ et al., 2018; Horvát et al., 2016; Markov et al., 2014; Oh et al., 2014; Rubinov et al., 2015; Scannell et al., 1995). However, a small proportion of strong long-range connections that deviate from the expectations of EDR model are consistently observed in empirical brain networks (Deco et al., 2021; Roberts et al., 2016). In addition, connectivity weights are typically lognormally distributed (Ercsey-Ravasz et al., 2013; Gămănuţ et al., 2018; Song et al., 2005; Wang et al., 2012), and nodal degrees are characterized by a scale-free distribution (Broido & Clauset, 2019; Eguiluz et al., 2005; Gastner & Ódor, 2016; Giacopelli et al., 2020; Sporns et al., 2004; van den Heuvel et al., 2008; Zucca et al., 2019). We examined whether our generative model recapitulates these properties. Figures 2 and 3 summarize the key findings in terms of variation in *β* and *L*_*s*_, respectively.

**Figure 2.**
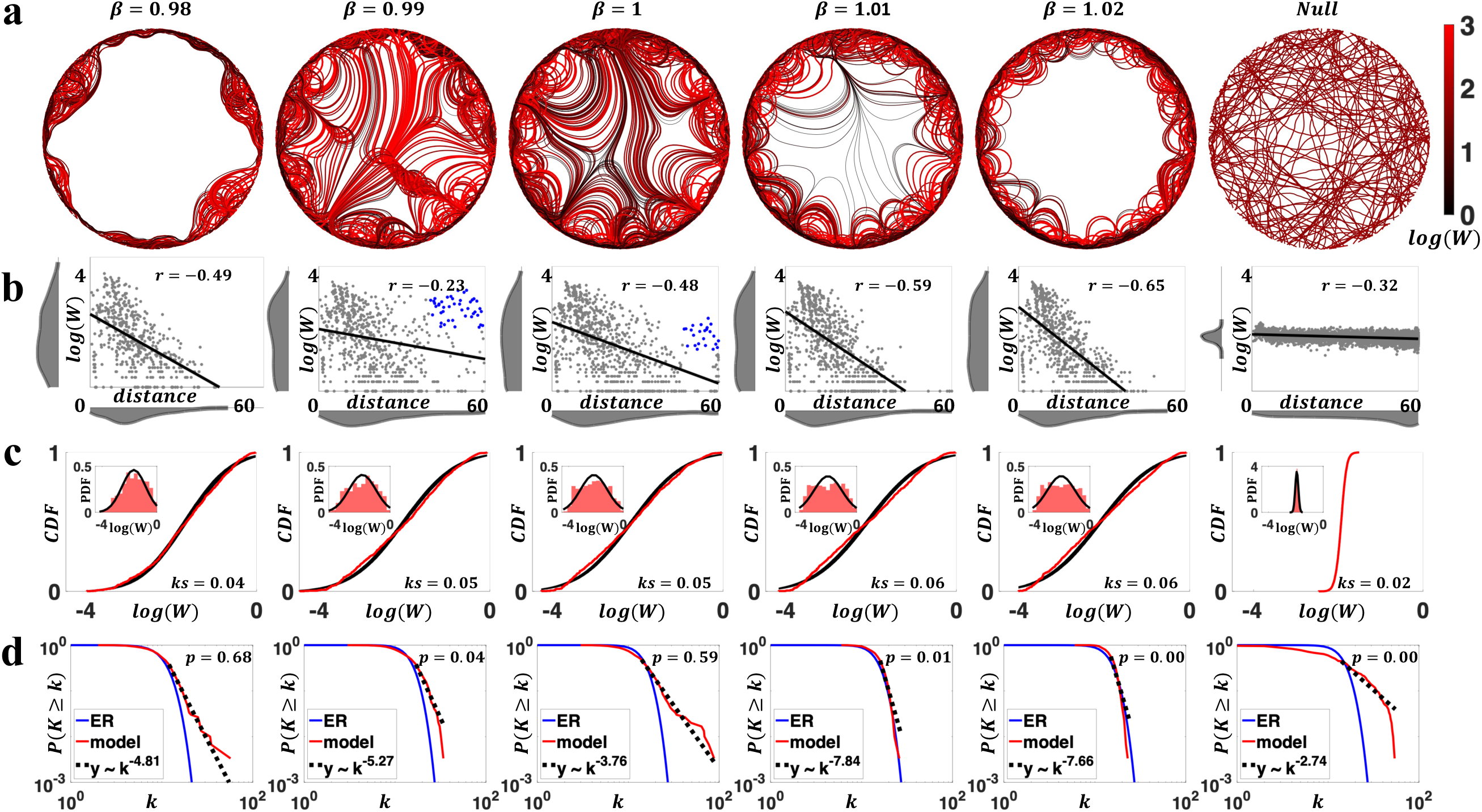
Characterization of generated connectomes under variation of the force decay parameter (i.e.,*s*). *L*_*s*_ was fixed to 1. **a)** Circles show generated axons for different values of *β* (0.98-1.02). Right most circle shows generated axons using a random walk null model. Axons are color-coded (using a black-red spectrum, see colorbar) by connection weights, such that black (red) curves represent weaker (stronger) connections. Color scale is truncated at a connectivity weight of 10^3^. The null network shows 5% of axons generated. **b)** Scatter plots of connection weights (log-scaled) versus distances for networks in panel a). Strong long-range connections that deviate from EDR are shown as blue dots in *β* = 0.99 and 1. Distributions of connection weights and distances are shown in marginal histograms. **c)** Distributions of connection weights (normalized by nodal strength) for networks in panel a) (red), compared to the fitted lognormal distributions (black), in terms of the cumulative density function (CDF, main figures) and probability density function (PDF, insets), respectively. KS described the one-sample Kolmogorov-Smirnov statistics of lognormal fit. **d)** Nodal degree distributions for evaluated *β* values. Results (with median *p*-value among 1,000 simulations; model and null) are compared to Erdös-Rényi random networks (ER) and scale-free fits (*y*∼*k*^−α^). Scale-free is plausible if *p* > 0.1.

**Figure 3.**
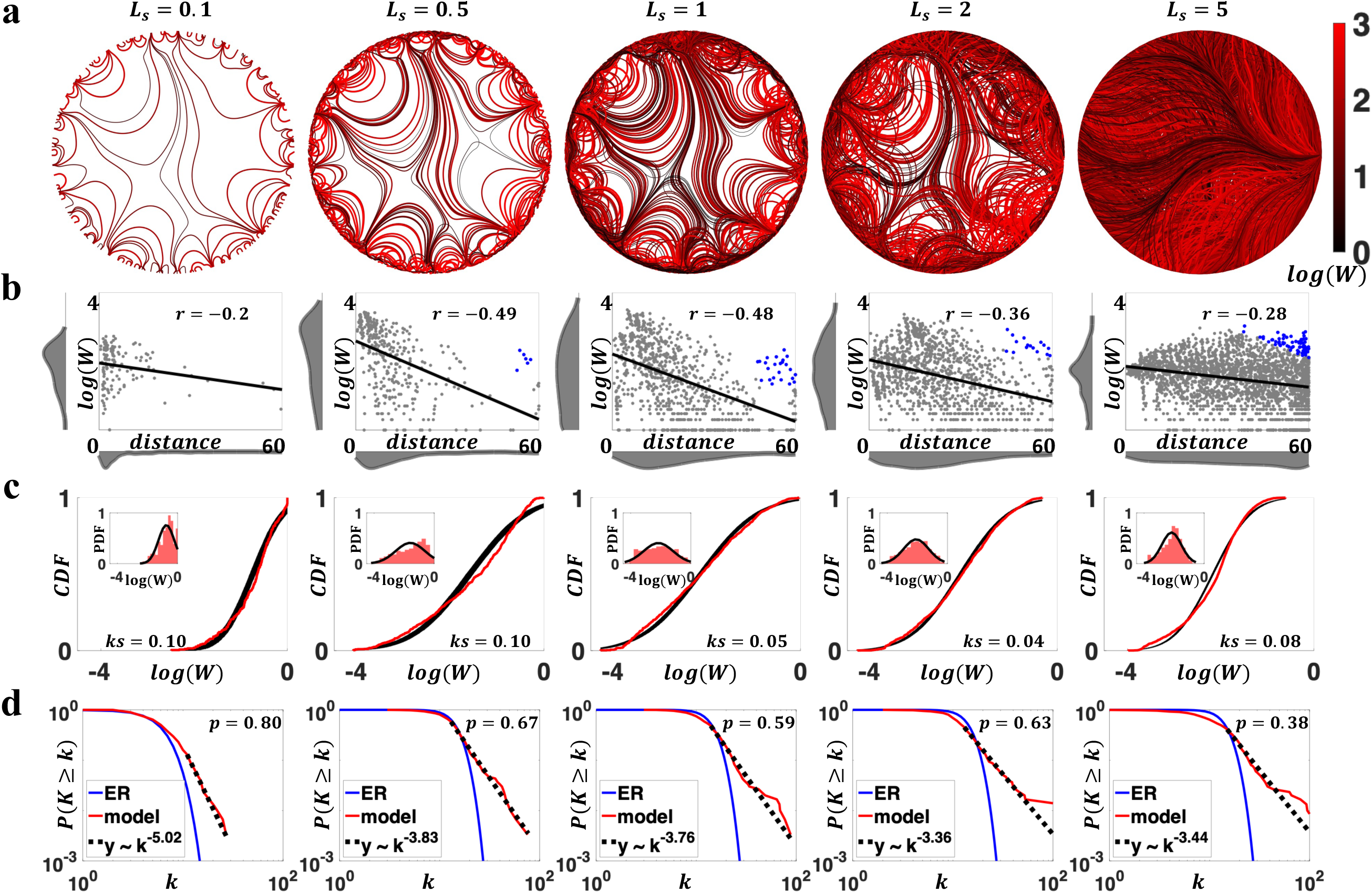
Characterization of generated networks changed under variation in the step length parameter (i.e., *L_s_*). Results are visualized for representative parameter combinations (*L*_*s*_ = 0.1 − 5 and fixed *β* = 1). **a)** Axon organizations of model networks. Higher network density was evident with increasing *L*_*s*_ (4,11, 24, 35, and 75% connectivity density, from left to right). **b)** Negative associations between connection weights and distances, with blue dots in *L*_*s*_ = 0.5, 1, 2 and 5 representing strong long-range connections that deviate from EDR. Distributions of connection weights and distances are shown in marginal histograms. **c)** Distributions of connectivity weights (normalized by nodal strength) in model networks (red), compared to fitted log-normal distribution (black). The main figure compared CDF, and the insets compared PDF. **d)** Degree distributions in generated networks, compared to ER networks and scale-free fit. All evaluated *L*_*s*_ values showed scale-free behaviors (median *p* > 0.1).

Figure 2 shows the effects of variations in *β*. We observed that small changes in *β* had marked effects in the topology of generated networks, especially in terms of the formation of long-range bundles of axonal projections. Specifically, when *β* was either small (*β* = 0.98) or large (*β* = 1.02), generated networks lacked distant connections (Fig 2a, 2b). This was because local nodal guidance was too weak (strong) to allow axon terminations (outgrowths) of long-range connections, as detailed in Fig. S2. Nevertheless, for moderate *β* values (*β* = 0.99, 1, and 1.01, Fig. 2a, 2b), strong long-range connections emerged, decreasing in strength with greater *β*. Remarkably, these connections replicated the strong long-range connections observed in empirical connectomes that cannot be explained by a simple EDR (Deco et al., 2021; Roberts et al., 2016). That is, it is not only a distance-dependence that matters, but the outgrowth of axons depending on competing attractive factors also contributes to brain connectivity profiles (Cahalane et al., 2011), including long-distance connections.

Compared to null networks generated from a constrained random walk (see Methods), our model networks featured brain-like axonal projections resembling U-fibers and white-matter tracts. As shown in Fig. 2a (most evident for *β* = 1), axonal projections were organized into bundles at a distance from their origins and defasciculated before reaching their targets. This is a natural consequence of axon guidance in the presence of target-released diffusible chemoattractant (i.e., the node exerted distance-dependent attractive force) that has been observed in past axon pathfinding studies (Hentschel & Ooyen, 1999). Adjacent nodes were connected via U-shape fibers that gradually steered according to the dynamically changing axon guidance. In contrast, axons generated by the null model failed to form organic fiber bundles. Additionally, the proportion of variance in model generated fiber length explained by Euclidean distance is comparable to empirically observed values (Akarca et al., 2021), a characteristic missing in the random walk null model (Fig. S4).

A negative association between connection weight and the Euclidean distance between nodes was evident across the range of *β* values considered (Fig. 2b). Connection weights (normalized by nodal strength, see Methods) spanned four orders of magnitude and were most parsimoniously modeled by lognormal distributions (Fig. 2c, Fig. S5). In contrast, connection weights for the random-walk null model were most accurately described by a gamma distribution (Fig. S5) and exhibited less variability, distinguishing them from model networks and empirical connectomes (Fig. 2b, 2c, Fig. S6).

To investigate whether our generated networks showed scale-free degree distributions, we adopted the approach developed by Clauset et al. (2009), as detailed in Methods. In brief, the approach returned a *p*-value describing the goodness-of-fit, and evidence of a scale-free distribution was deemed plausible if more than 50% of the networks in a population had a *p* > 0.1 (Broido & Clauset, 2019). For each representative parameter combination considered, we generated 1,000 networks and evaluated evidence for scale-free degree distributions. Fig. 2d shows the networks with the median *p*-value for each parameter combination (see Fig. S7 for *p*-value distributions). Scale-free behavior was evident for *β* = 0.98 and 1, yet it was missing in networks generated with other *β* values and our null networks. Crucially, *β* = 1 generated networks that were simultaneously characterized by all the connectomic properties evaluated in this section.

We next characterized the impact of variation in step length, *L*_*s*_, on properties of the generated connectomes. Greater *L*_*s*_ led to networks with higher connection density (ranging from 4% - 75% in Fig. 3a). This was because a larger *L*_*s*_ resulted in fewer trajectory updates, and thus the past guidance an axon received had a longer-lasting impact on subsequent wiring. As a result, if two axons originated from the same node yet different coordinates, with a greater *L*_*s*_, the difference in initial coordinates made their subsequent trajectories less likely to converge, which led to more diverse wiring and denser networks. We also found that the choice of *L*_*s*_ impacted connectivity weight distributions (Fig. 3b, 3c). Certain values of *L*_*s*_ (*L*_*s*_ = 1 and 2, Fig. S5) generated networks with lognormally distributed connectivity weights. Negative associations between connection weights and distances were consistently observed, and strong long-range connections that deviate from EDR were found in generated networks except for *L*_*s*_ = 0.1 (Fig. 3b). In addition, all representative *L*_*s*_ parameters generated networks with scale-free degree distributions (Fig. 3d, S8).

Collectively, these results suggested that combinations of *β* and *L*_*s*_ generate connectomes with realistic properties, including scale-free degree distributions, brain-like axonal bundles, negatively correlated connection weight and distance, log-normally distributed weights, and strong long-range connections that deviate from EDR. Specifically, comparing the two model parameters *β* and *L*_*s*_, variations in *β* had a stronger effect on degree distributions, whereas changes in *L*_*s*_ were more closely related to weight distributions.

### Emergence of complex network properties

So far, we have focused on characterizing the distributions of connection weight and nodal degree generated by our model. In this section, we examine whether our generative model could give rise to connectomes exhibiting complex topological properties, including small-worldness and modularity. We specifically investigated the network average clustering coefficient (CC), characteristic path length (CPL), small-worldness (SW), and modularity Q, benchmarked to weight and degree preserved null networks (see Methods). These topological features are hypothesized to relate to the functional segregation and integration of brain networks (Bassett & Bullmore, 2017; Fornito et al., 2015; Sporns & Betzel, 2016).

Using an exhaustive grid search (see Methods), we investigated how weighted topological properties of generated networks changed as a function of *β* and *L*_*s*_. Fig. 4a displays the variation of network topology among the parameter space that generated brain-like connection weight and nodal degree distributions. Topological properties were evaluated at a network density of 10%, yet the patterns of topological variations were insensitive to network densities (Fig. S8). Across the investigated parameter space, generated networks consistently displayed small-world and modular structures. A higher CC and a longer CPL were evident relative to weight and degree preserved null networks. Variations in model parameters impacted the network topology in a continuous manner. Increasing *β* and decreasing *L*_*s*_ led to weighted networks with higher clustering, longer characteristic path length, stronger small-worldness, and weaker modularity (also see Fig. S9).

**Figure 4.**
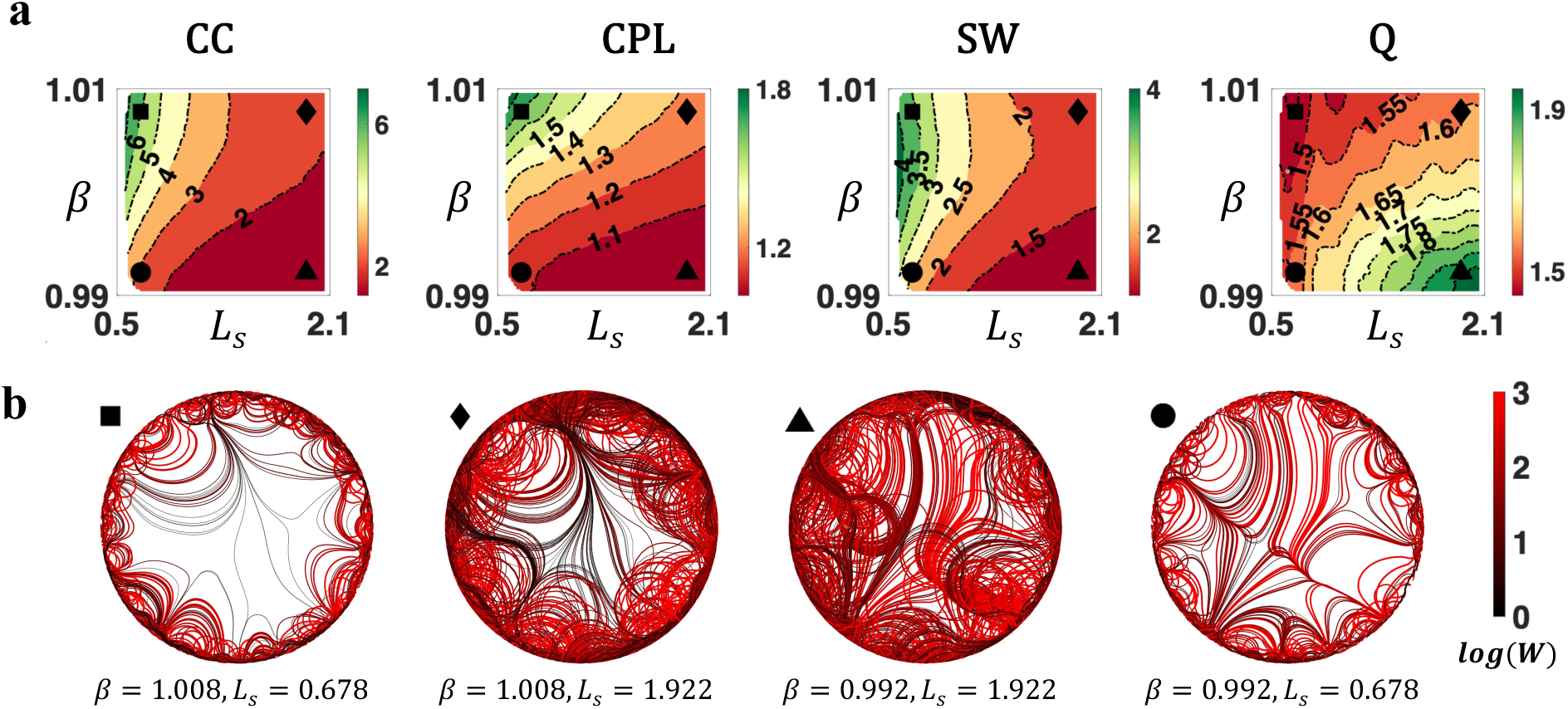
Complex topological organization of the generated connectomes. **a)** Contour plots of weighted network average clustering coefficient (CC), characteristic path length (CPL), small-worldness (SW), and modularity Q (Q) of generated networks, benchmarked to null networks with preserved weight, degree, and strength distributions. All measures were normalized to the null networks. **b)** Axon organization of example networks, generated by parameters labelled with square (*β* = 1.008, *L*_*s*_ = 0.678), diamond (*β* = 1.008, *L*_*s*_ = 1.922), triangle (*β* = 0.992, *L*_*s*_ = 1.922), and circle (*β* = 0.992, *L*_*s*_ = 0.678) in Fig. 4a.

To supplement this analysis, we visualized the axon organizations of example networks from different positions of the parameter space. As shown Fig. 4b, decrease in *β* and increase in *L*_*s*_ improved the prevalence and strength of medium-to-long range connections, reducing segregation (measured by clustering) while promoting integration (measured by efficiency, i.e., the inverse of characteristic path length).

In summary, variations in model parameters shifted network topology by adjusting the balance between short-range and long-range connectivity. While the generated networks consistently showed small-world and modular organizations, the degrees of these properties vary. Combined with the results from Fig. 2 and 3 (i.e., analyses on bundle structure, connection weights, and degree distributions), certain combinations of model parameters (e.g., *β* = 1 and *L*_*s*_ = 1) were capable of generating networks that resembled all the evaluated connectomic features.

### Generating connectomes for individuals

Finally, we focused on fitting the two parameters governing our generative model to individual human connectomes. Our motivations here were two-fold. First, the results above suggested networks generated by our model exhibited varying degrees of small-worldness and modularity. It was unclear if certain parameters gave rise to these properties numerically close to empirical connectomes, and if these parameters resided in the range that generated networks simultaneously manifested brain-like connection weights, nodal degrees, and axonal bundles. This question could be answered via a parameter optimization approach. Second, model optimization enables comparison of individuals in terms of their fitted parameters and can provide insight into inter-individual variability in the processes guiding connectome development. For example, inter-individual variation in parameters of existing generative models is associated with various traits, including age, sex, a schizophrenia diagnosis, and social-economic disadvantage (Akarca et al., 2021; Betzel et al., 2016; Carozza et al., 2023; Faskowitz et al., 2018; Simpson et al., 2011; Sinke et al., 2016; Siugzdaite et al., 2022; Zhang et al., 2021). Thus, a parameter fitting scheme is a prerequisite for future cohort studies to apply our model.

Parameters for the current state-of-the-art connectome generative models are typically optimized by minimizing an energy function that compares the degree, clustering, betweenness centrality, and Euclidean space edge lengths between generated and empirical connectomes (Betzel et al., 2016). However, our model does not generate networks with nodes that correspond one-to-one to empirical connectomes; as a result, the discrepancy in Euclidean space edge lengths cannot be evaluated and the classical energy cost is not applicable (despite this, when evaluated using the degree, clustering, and betweenness centrality, our model achieved a better fit relative to the established geometric model; see Fig. S10). Therefore, we developed a new method to fit the parameters of our generative model to individual connectomes (detailed in Methods) and applied it to estimate parameters for 1,064 participants in the Human Connectome Project (HCP) (Van Essen et al., 2013).

The parameter estimation method aimed to minimize the dissimilarity between the generated network and an individual’s connectome, in terms of CC, CPL, and modularity Q. Fig. 5a displays the best-fit model parameters for the HCP population, and networks generated by the fitted parameters numerically resembled topological features for which they were optimized (Fig. S11). Individual parameters were found significantly different between males and females (*p* = 2*e*^*&+^, Fig. S12), confirming the model’s capability of capturing inter-individual variations in connectomes. Using the group averaged model parameters (*β* = 0.9968, *L*_*s*_ = 0.9531), we generated networks and investigated their weight, degree, and axonal bundle structures. As shown in Fig. 5b, generated networks simultaneously manifested brain-like axonal bundles, negatively correlated connection weight and distance, log-normally distributed weights, scale-free degree distributions, and strong long-range connections that deviate from EDR. These results suggested a “sweet spot” of parameters can be identified to generate human brain-like connectomes possessing a host of empirically observed properties.

**Figure 5.**
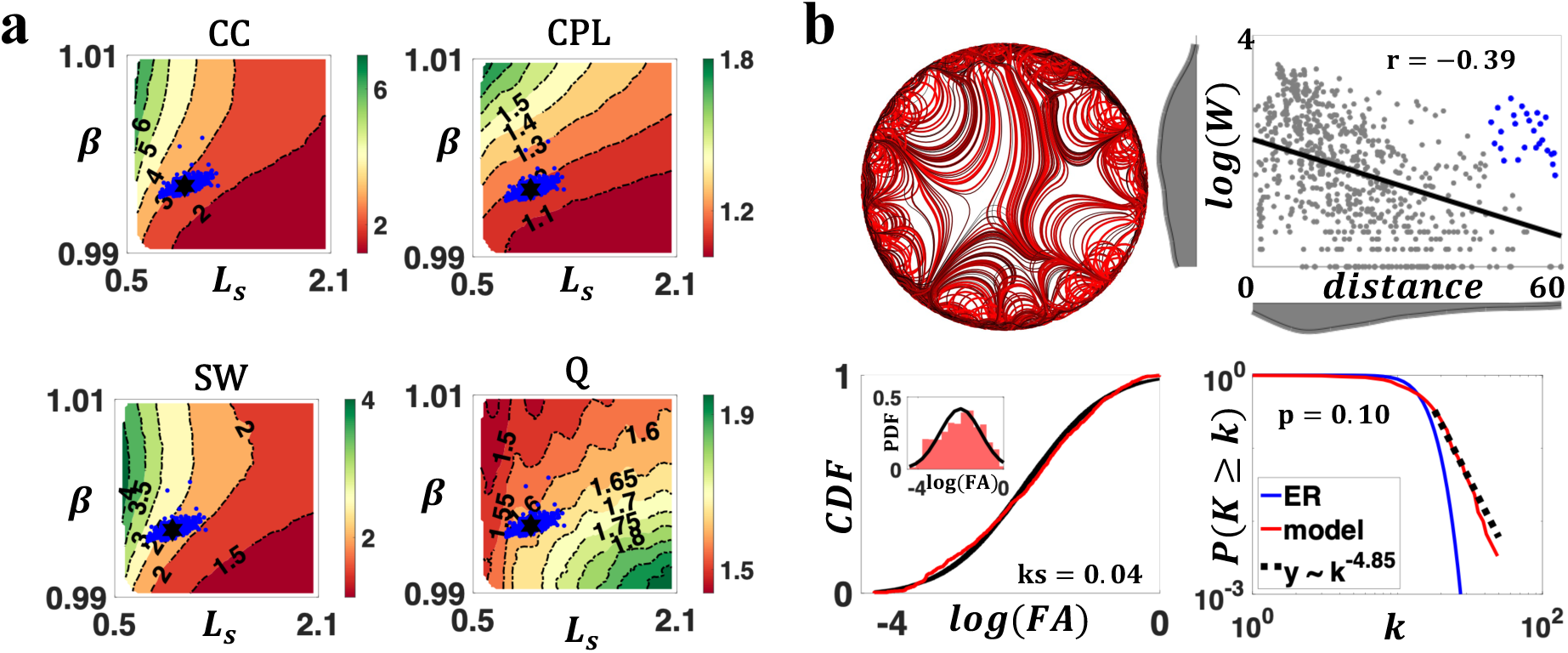
Individual parameters for HCP connectomes. **a)** Optimized individual parameters (blue dots) overlaid on contour plots of weighted topological measures (CC, CPL, SW, and modularity Q). The black hexagram represents the group average parameters (*β* = 0.9968, *L*_*s*_ = 0.9531). **b)** Networks simulated with the HCP group average parameters showed organic axon organization (top left), negatively associated connection weights and distances (top right; blue dots represent strong long-range connections that deviate from EDR), lognormally distributed weights (bottom left), and scale-free degree distributions (bottom right, despite the *p* value marginally above the threshold of *p* = 0.1).

## Discussion

Generative models can provide insight into the wiring principles governing brain network organization. Most existing models generate unweighted connectomes in which connections are either present or absent (Betzel et al., 2016; Oldham et al., 2022; Simpson et al., 2011; Sinke et al., 2016; Vértes et al., 2012). However, it is well-known that connectivity weights are diverse and span multiple orders of magnitude (Fornito et al., 2016). In this work, we developed a novel model that generates connectomes weighted by axon counts and demonstrated that our model can recapitulate lognormal connectivity weight distributions. Our model is spatially embedded and characterizes the dynamics of axonal outgrowth. We demonstrated that our model manifests key topological properties of the connectome and yields axonal fiber bundle structures that resemble white matter fascicles. We were also able to fit our generative model to individual connectomes, enabling future cohort studies to apply the model.

Our work substantially builds on the seminal generative model developed by Song et al. (2014). Whereas this earlier model considered static axon propagation in a fixed direction, we established a dynamic axon pathfinding model in which the attractive forces guiding axonal outgrowth are continuously updated. With appropriate selection of *β* and *L*_*s*_, we observed that our model can generate networks with properties that are consistent with empirically mapped connectomes. In particular, negative associations between connectivity weights and the Euclidean distances between nodes were evident—a property that has been found in numerous studies (Betzel & Bassett, 2018; Ercsey-Ravasz et al., 2013; Roberts et al., 2016). We also found that connectivity weights were lognormally distributed and showed scale-free degree distributions. This suggests that a simple axon growth mechanism can generate key properties of a connectome’s topological architecture.

The choice of *β* and *L*_*s*_ determines whether the generated networks show complex topologies. As *β* is increased, distant nodes exert less influence on an axon’s growth and thus axons are attracted by neighboring nodes, forming short-range connections. This leads to the formation of clusters between spatially adjacent nodes and weakens the long-range connections that are critical to network efficiency. In contrast, a larger step length parameter *L*_*s*_ enables axons to propagate beyond a local nodal sphere of influence, increasing the prevalence and strength of long-range connectivity and improving network integration. While *β* and *L*_*s*_ impact generated axons through mechanisms that are both similar and distinct, their combined effects lead to generated networks with small-worldness and modularity, akin to empirical connectomes.

Biological plausibility is a key characteristic of our model, where axon outgrowth is determined by the summed attractive forces exerted by each node. This is inspired by the observation that axons grow by responding to complex and combined effects of multiple guiding cues (Wadsworth, 2015). The two governing parameters also build on empirical observations in neural systems. The force decay parameter *β* determines the distance-dependent decay of attractive forces exerted by nodes, modeling the concentration decay of guiding cues with distance from releasing sites (Kaiser et al., 2009; Murray, 2002). The step length parameter *L*_*s*_ governs the extent to which an axon can change its trajectory per unit length, and it can represent the combined effects of multiple factors such as a growth cone’s growing speed and its sensitivity to molecular guidance (Alberts, 2017). While step length is seldom considered as a key parameter in applications such as tractography (Tournier et al., 2002), *in vitro* evidence suggests it is a vital factor that models axonal growth. For example, variations in growth step length have been observed between frog and chick neurons, as well as between normal and regenerating frog neurons (Katz et al., 1984).

Our model generates connection weights by counting axons between nodes, a method distinct from other recently proposed models. Different weight inference approaches all have their unique strengths. Using a weighted stochastic block model, Faskowitz and colleagues (2018) inferred connection weights from network blocks, highlighting the community architecture of connectomes. Based on an unweighted connectome generative model, Akarca and colleagues (2023) introduced connection weights via minimizing the redundancy in network communicability, capturing dynamics in the strengthening and weakening of connections. In contrast, axon counts used in our model are intrinsically akin to streamline counts synonymous with structural connectomes, emphasizing the physical nature of connections as neural pathways.

Elucidating the mechanisms governing the formation of long-range connections remains a pivotal yet unsolved question in connectome generative model research. Early work suggested that, in addition to the distance rule, a topological homophily rule is required to promote the formation of long-range connections (Betzel et al., 2016; Vértes et al., 2012). Recently, the biological plausibility of topological rules was questioned, and homophily in gene expression and cytoarchitecture was hypothesized to contribute to long-range wiring (Kerstjens et al., 2022; Oldham et al., 2022). Nevertheless, existing frameworks failed to explain the specificity of long-range connectivity (Betzel & Bassett, 2018). Moreover, it is unclear how brain elements can perceive distant pairs without prior global knowledge of topology. Due to the spatial embedding of brain networks, a distance component might be required for brain elements to search for their wiring pairs. By including a pathfinding component, our model simulated long-range connections, including those that deviate from the EDR. Growth cones were sequentially guided by the strong local cues of a series of intermediate nodes before reaching their distant destinations (despite weak attractive forces exerted by distant nodes still contribute). This mechanism is consistent with the hypothesis of intermediate targets in axon guidance, whose suggestive evidence has been observed in model organisms such as *Drosophila* and mice (Canty & Murphy, 2008; Dickson, 2002).

We conclude by acknowledging the limitations of our work and providing guidance for future improvement. Firstly, as the first attempt to generate connectomes from dynamic axon guidance, the model simplifies the brain as a two-dimensional circle and ignores complex brain structures such as sulci, gyri, deep gray matter, and cerebrospinal fluid. While this approach contributed to model simplicity and axon visualization, it also introduced limitations, such as the loss of nodal correspondence between generated and empirical connectomes. Using a realistic brain mesh/volume to incorporate three-dimensional neuroanatomical constraints in axonal outgrowth would naturally address these limitations but also entail higher computational demands. Secondly, we assume that all brain regions have the same distance-dependent attractiveness, and that all axons are equally sensitive to guidance from brain regions. These assumptions are likely breached in the brain given the diversity in regional properties (e.g., cortical thickness, curvature of folds, laminar structure, cellular composition, and neuronal density), neuron types, and guiding cues (attractive and repulsive, chemical and mechanical). Recent efforts in generating high-resolution brain maps such as molecular and cytoarchitectural profiles (Amunts et al., 2013; Arnatkevičiūtė et al., 2019; Hansen et al., 2022; Markello et al., 2022) might provide an opportunity to refine the assumptions and improve the model’s capacity. Thirdly, while our model generates axon organization that is visually akin to axon bundles and white matter fascicles, factors that contribute to axon bundling are not considered. Incorporating fasciculation mechanisms such as the contact attraction between axons and axon-released guiding cues (Hentschel & Van Ooyen, 2000) might help to build a more nuanced white matter and connectome organization. In addition, the model implements a deterministic axonal guidance rule, and as such, stochasticity, which is also fundamental to neural development (Carozza et al., 2023; Hassan & Hiesinger, 2015), was not taken into account. Future work could evaluate the robustness of the model with the presence of stochasticity, such as random noise in guiding cues and axon growth. Finally, in this study, our model generates macroscale connectomes, yet this is achieved by simulating axons that are microscale anatomical concepts. Future studies could investigate our model’s application in generating microscale connectomes.

## Acknowledgements

Data were provided by the Human Connectome Project, WU-Minn Consortium (Principal Investigators: David Van Essen and Kamil Ugurbil; 1U54MH091657) funded by the 16 NIH Institutes and Centers that support the NIH Blueprint for Neuroscience Research; and by the McDonnell Center for Systems Neuroscience at Washington University. The computation is supported by The University of Melbourne’s Research Computing Services and the Petascale Campus Initiative. Y.L. is funded by the Melbourne Research Scholarship. C.S. is supported by the Australian Research Council (grant number DP170101815). R.F.B is supported by the National Science Foundation (award ID: 2023985). D.A. is supported by the James S. McDonnell Foundation Opportunity Award and the Templeton World Charity Foundation, Inc. (funder DOI 501100011730) under the grant TWCF-2022-30510. M.A.D. was supported by an NHMRC Investigator Grant (1175754). A.Z. is supported by research fellowships from the NHMRC (APP1118153).

## Code and data availability

The data of HCP is publicly available from https://www.humanconnectome.org/. The code for the generative model is available from https://github.com/yuanzhel94/connectome_from_pathfinding.

## Methods

### Model implementation

An overview of our model is described in the Results section. Here, we provide finer details of the model, elaborating on aspects including node heterogeneity, path constraints, axon termination, and parameter specifications.

To parcellate the hypothetical gray matter, *N*_*n*_ node centers were evenly positioned along the circle perimeter, such that the angular distance between adjacent nodes equals 2*π*/*N*_*n*_. Next, nodal heterogeneity was introduced by randomly perturbing node center coordinates. This was accomplished by applying a uniformly distributed angular displacement, *ε*∼*ρ* ∗ (−*π*/*N*_*n*_, *π*/*N*_*n*_), to each node center. Specifically, *ρ* = 1 was used in this study to maximize nodal heterogeneity while preserving the sequential arrangement of nodes along the perimeter.

Axons were simulated based on the distance rule in *Eq*. 1. To encourage axons to traverse relatively non-curved trajectories, regularity constraints were applied to each axon from the second extending step onward. The regularity constraints stipulate that the angle formed between the direction of two consecutive steps cannot exceed the angle *θ*. In other words, if the angle between two consecutive steps exceeds *θ*, the second step is adjusted such that the angular difference is forced to *θ* (Fig. S1).

Ideally, axons would terminate on the circle circumference, connecting two points of the hypothetical gray matter. However, not all simulated axons can successfully reach the circle perimeter. When the value of *β* was small, a “black hole” region emerged within the circle, as shown in Fig. S2. Axons entering the “black hole” cannot escape, forming a circular trajectory of infinite loops. To address the problem, a parameter *S*_*max*_ was introduced to stipulate the maximum number of growing steps allowed. Axons failing to reach the circle circumference within *S*_*max*_ steps were considered unsuccessful and were excluded from network construction and analyses.

Eight parameters were defined in the model. Unless otherwise specified, default values of parameters (Table 1) were used. A comprehensive justification for parameter choice was included in Supplementary Materials.

**Table 1.** Default values of model parameters.

| Parameters | Meaning explained | Default values |
| --- | --- | --- |
| $R$ | Circle radius | 30 |
| $N_n$ | Number of nodes | 84 |
| $\rho$ | Controls nodal heterogeneity | 1 |
| $N_a$ | Number of axons | $2e^5$ |
| $\theta$ | Angular constraint | $15^\circ$ |
| $\beta$ | Power-law decay of attractive force | To be optimized |
| $L_s$ | Growth step length | To be optimized |
| $S_{max}$ | Maximum growing steps | $3R/L_s$ |

### Weight and degree measures of generated networks

We investigated the associations between edge weights and distances, and the weight distributions in generated networks. The weight-distance associations were evaluated by calculating the Pearson’s correlation coefficient between edge lengths (i.e., the Euclidean space distance between two nodes connected by an edge) and the common logarithms of the edge weights. The weight distributions were also described in the common logarithm scales; however, instead of using the raw weights (*C_ij_*), weights normalized by nodal strengths (*A_ij_* = *C_ij_*/ ∑_*k*_ *C_ik_*) were utilized. These normalized weights quantified the fraction of axons maintained by node *j* that connected to node *i*, conceptually replicating the fraction of labeled neurons in Ercsey-Ravasz et al. (2013) that was found lognormal. Weight distributions were evaluated against fitted lognormal, gamma, normal, exponential, and Weibull distributions using one-sample KS test.

We also analyzed the degree distributions of generated networks. To reduce the bias of finite network size, 1,000 networks, each comprising 300 nodes (*N*_*n*_ = 300), were generated for each evaluated parameter combination. Next, generated networks were threshold and binarized to a network density of 5% (except *β* = 0.98, 0.99, and *L*_*s*_ = 0.1 that were evaluated at a lower density because their generated networks are too sparse. However, these parameters do not generate brain-like networks). To assess the scale-free property of degree distributions, we employed the method developed by Clauset et al. (2009). Consider a network whose nodal degrees *K* adhere to a scale-free distribution for *K* ≥ *K_min_*, its probability density function is given by

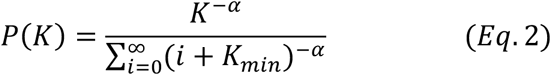

The Clauset method estimated *K_min_* by a Kolmogorov-Smirnov minimization approach and optimized *α* through a maximum likelihood estimation. The goodness-of-fit was assessed with a bootstrap approach, and the null hypothesis of scale-free was rejected if *p* < 0.1. Applied to a network population (in our study, 1,000 networks generated from the same model parameters), scale-free was deemed a plausible hypothesis if more than 50% networks showed *p* ≥ 0.1. Further details of the scale-free test can be found in Broido and Clauset (2019).

Results of weight and degree analyses were visualized for representative parameters (*L*_*s*_ = 1, *β* = 0.98, 0.99, 1, 1.01, and 1.02; *β* = 1, *L*_*s*_ = 0.1, 0.5, 1, 2 and 5). These parameters were selected to generate diverse network properties while delineating the isolated effects of each parameter. Compared to an exhaustive grid search (used in a later section to evaluate global topology), this approach enabled us to uncover details (Fig. 2 and 3) that were obscured in summary metrics (i.e., Pearson *r*, KS statistics, and *p*-values).

Null networks generated from a constrained random walk were used to benchmark model networks. Specifically, axon growth directions were randomly sampled from (−*θ*, *θ*) rather than being calculated from the distance rule in Eq. 1. Step length parameter of *L*_*s*_ = 1 was used. All other parameters remained consistent with the model.

### Global topology of generated networks

To characterize the global topology of generated networks, model parameters were drawn from a grid combination of *β* and *L*_*s*_ (0.99≤ *β* ≤ 1.01, 0.1≤ *L*_*s*_ ≤ 2.1; 101-by-101 grid). This parameter space was determined from preliminary experiments and was found to generate networks that replicated connectomic features. To account for the stochastic variability arising from node and axon sampling, fifty networks were generated for each parameter combination, forming 50 network landscapes. The network topology corresponding to each parameter combination was described by the average topological metrics over 50 landscapes.

We considered the weighted clustering coefficient, characteristic pathlength, small-worldness, and modularity Q of generated networks. Because topological measures are fundamentally related to network density and connectivity strengths, all generated networks were threshold and normalized to have the same network density (10%) and total connectivity (2*e*^+^). Parameters whose generated networks have a density smaller than 10% were ignored. Topological measures were evaluated using the Brain Connectivity Toolbox (BCT), benchmarked to weight and degree preserved null networks constructed using the null_model_und_sign() function in BCT.

### Empirical datasets

This study utilized the Human Connectome Project Young Adults (HCP, 1064 subjects) datasets (Glasser et al., 2013; Uğurbil et al., 2013). A comprehensive description of data acquisition and connectome construction has been detailed elsewhere (Mansour L et al., 2021). The HCP connectomes were mapped to the Desikan-Killiany atlas, comprising 68 cortical and 16 subcortical brain regions. Networks were threshold to a density of 10%.

### Optimize model parameters against connectomes

We optimized the model parameters for the HCP connectomes. Because topological measures are related to network density and connectivity strengths, empirical and model networks were threshold and linearly scaled to the same network density and total connectivity (discussed in supplementary materials). Parameters were fitted to minimize the discrepancies between empirical and model networks, measured by the rooted mean squared error (RMSE) in weighted CC, CPL, and modularity Q (Eq. 3). Small-worldness was excluded because it is a combination of CC and CPL.

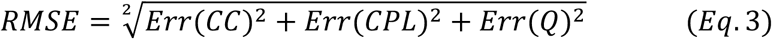

To mitigate the inconsistent scales among topological measures, metrics were normalized by the values in degree and strength preserved null networks and standardized using the standard deviation in empirical connectomes.

Parameters were optimized using a Monte Carlo method through an exhaustive grid search (see Methods: Global topology of generated networks). To account for the stochasticity-dependent inaccuracy and unreliability, and to improve the computational tractability, we employed the fast landscape generation (FLaG, generating 50 landscapes) and the multilandscape method developed by Liu et al. (2023). For each landscape, the best-fit parameters (with the smallest RMSE, values shown in Fig. S11) were selected, and the average across 50 landscapes was deemed the optimal parameters.

