## Supplementary for "A generative model of the connectome with dynamic axon growth"

The model specified the maximum angular disparity $\theta$ an axon can make at each increment step. Fig. S1 illustrates how this angular constraint is applied.

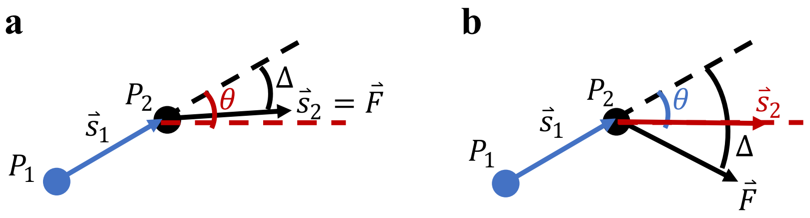


**Figure S1. Angular constraint $\boldsymbol{\theta}$.** *a) axons grow in the direction of the summed force if the computed angular difference is smaller than or equal to $\theta$. In growth step $i-1$, the axon grew from point P1 (blue dot) to point P2 (black dot), and its growth course in step $i-1$ is $\vec{S}_1$ (blue arrow). The black arrow denoted the summed force $\vec{F}$ that the axon received at position P2. $\Delta$ represented the angular difference between $\vec{F}$ and $\vec{S}_1$, and the angle $\theta$ between black and red dashed lines was the maximum angular disparity allowed. When $\Delta \leq \theta$, the axon's growth direction in step i ($\vec{S}_2$) equals $\vec{F}$. b) Same as a) but for $\Delta > \theta$, the angular difference between $\vec{S}_1$ and $\vec{S}_2$ (red arrow) is forced to $\theta$.*

In addition, two special cases were considered.

Case 1: if $\vec{\mathrm{F}}$ and $\vec{s}_1$ are in opposite direction ($\frac{\vec{\mathrm{F}}}{|\vec{F}|} + \frac{\vec{s}_1}{|\vec{s}_1|} = 0$), the clockwise/counterclockwise direction of $\theta$ could not be determined. In this scenario, we forced $\vec{s}_2$ to be in the direction of $\vec{s}_1$.

Case 2: if $\vec{F}$ has a magnitude of 0, its direction cannot be determined. In this scenario, the model randomly sampled a non-zero $\vec{F}$, after which the angular constraint $\theta$ was applied.

***"Black hole" region and $\boldsymbol{\beta}$ values***

In Results and Methods, we argued that growing axons could fail to land on the circle circumference because they were trapped within a “black hole” region. This happened when the values of $\beta$ were small, such that the guidance from local nodes was too weak to allow axon terminations. As shown in Fig. S2, as $\beta$ increases, trapped axons suddenly escape to form middle-range to long-range connections. However, if $\beta$ is too large, long-range connections cannot form because of the strong attractive force from local nodes. This explains why small variations in $\beta$ leads to dramatic change in generated networks: only a small critical range of $\beta$ allows long-range connections to form.

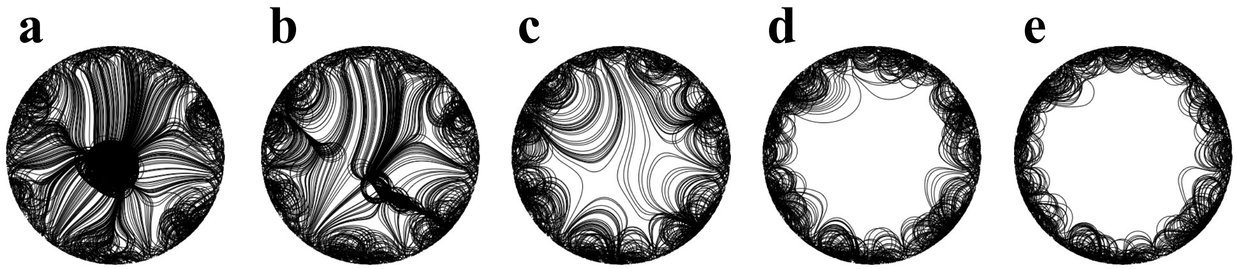


**Figure S2. Effect of $\beta$ on axon length.** *a)-e) each displayed a subset of 1,000 simulated axons from networks shown in Fig. 2a. $\beta = 0.98, 0.99, 1, 1.01$, and, $1.02$ respectively (from a to e). a) A “black hole” region is evident with small values of $\beta$. Axons were trapped and failed to form long-range connections. b)-c) As $\beta$ increases, long-range connections start to appear. d)-e) If $\beta$ is too large, long-range connections disappear due to strong local attractive forces.*

#### ***Parameter specification***

Eight model parameters were defined in the model. In this study, we were interested in variations in $\beta$ and $L_s$; other parameters were fixed. Here we provided a brief justification to this implementation.

$N_n$ (the number of nodes) is determined by the data of interest. In this study, we used 84 nodes to correspond to the number of brain regions in the Desikan-Killiany atlas. In addition, 300 nodes were used in scale-free analysis to reduce the bias from finite network size.

The choice of $\rho$ (controlling the node heterogeneity) has been explained in Methods. $\rho = 1$ was used in this study to maximize nodal heterogeneity while preserving the sequential arrangement of nodes along the perimeter.

$N_a$ (the number of axons simulated) was fixed to $2e^5$. This value was chosen as a balance between the favor to large $N_a$ and the computational affordability. The axon origins were sampled in a random manner; thus, a large $N_a$ is favored to capture the connectivity distribution of generated networks. Consider two networks (connectivity matrices $M_1$ and $M_2$) that only differ in the number of axons simulated ($N_a$ and $cN_a$, respectively; $N_a \to \infty$, $c$ is a constant). The connectivity expectation would be $M_2 = cM_1$. This further guided the trick (linear scaling) used in parameter optimizations (See Parameter optimization section in supplementary materials).

The effects of R (circle radius), $\theta$ (angular constraint), and $L_s$ (growth step length) on generated networks were closely related. Fig. S3 demonstrated a linear relationship between R and $L_s$. The association between $\theta$ and $L_s$ is complex and non-linear; however, the ratio $\theta / L_s$ could be approximated as the angular changes that an axon can make per unit growth length. Therefore, we decided to fix R and $\theta$ and investigate the variations in $L_s$, making $L_s$ a key tunable parameter in the model. R was set to a random constant ($R = 30$ was used), and the choice of $\theta = 15°$ was inspired by the angle values typically used between successive steps in tractography.

$S_{max}$ (the maximum number of growing steps allowed) ensured that simulations could stop even if axons were trapped by the “black hole” zone. $S_{max} = 3R/L_s$ was used such that axons were able to connect furthest points on the circle (Euclidean distance of 2R), while a margin of R was included to allow curved axon trajectories.

$\beta$ describes the distance decay of attractive force, controlling the relative contribution of guidance exerted by adjacent and distance nodes. As a result, values of $\beta$ are closely related to topology of generated networks, and we make it a key tunable parameter of the model.

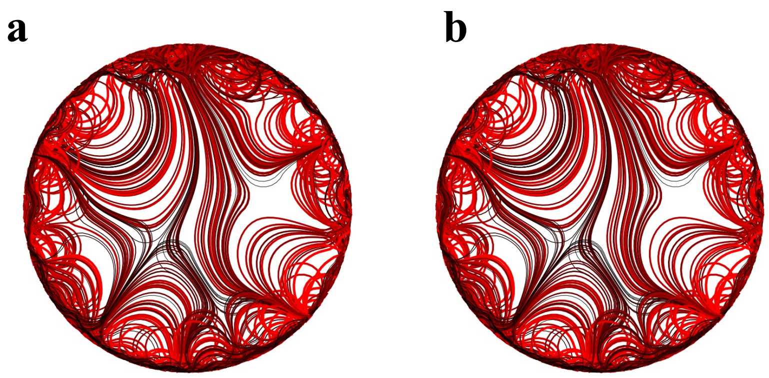


***Figure S3. R and $L_s$ are linearly related.*** *a) A network generated with R = 30 and $L_s = 1$. b) A network generated with R = 60 and $L_s = 2$. All other parameters, including the angular coordinates of nodes and axon origins, were the same. The two parameter combinations generated the same network.*

#### *Variance in fiber lengths explained by Euclidean distances*

Previous work found that the variance in fiber lengths explained by Euclidean distances is inconsistent between studies, ranging from 22-79%. Despite the wide range reported, the results from our model generated axons fell within the range (generally between 70-80%), whereas values from the random walk null model generated axons were much larger (Fig. S4).

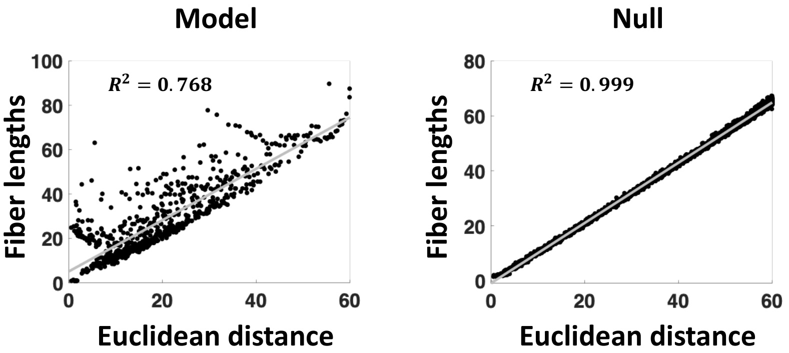


*Figure S4. Variance in fiber lengths explained by Euclidean distances is consistent with empirical observations in model but not in null.*

#### *Weight distributions (null model, and alternative distributions ks test results)*

In the main text, using representative parameter combinations, we suggested that the model was able to simulate networks whose connection weights were best described as lognormal distributions. Here, we statistically compared the fitted KS statistics among candidate distributions (lognormal, normal, gamma, exponential, Weibull, Fig. S5). 50 networks were generated for each parameter combination, and the KS statistic of fit were compared between distributions using paired t-tests. Lognormal distribution was found to be the best fit (significantly smaller KS statistic compared to other distributions, all $p < 0.05$) in networks generated with $\beta = 0.98, 0.99, 1, 1.01$, and $1.02$ ($L_s$ was fixed to 1), and $L_s = 1$ and 2 ($\beta$ was fixed to 1). It was also the best fit in networks generated with the group average parameter for the HCP population, despite the KS statistic difference between lognormal and Weibull distributions were not significant.

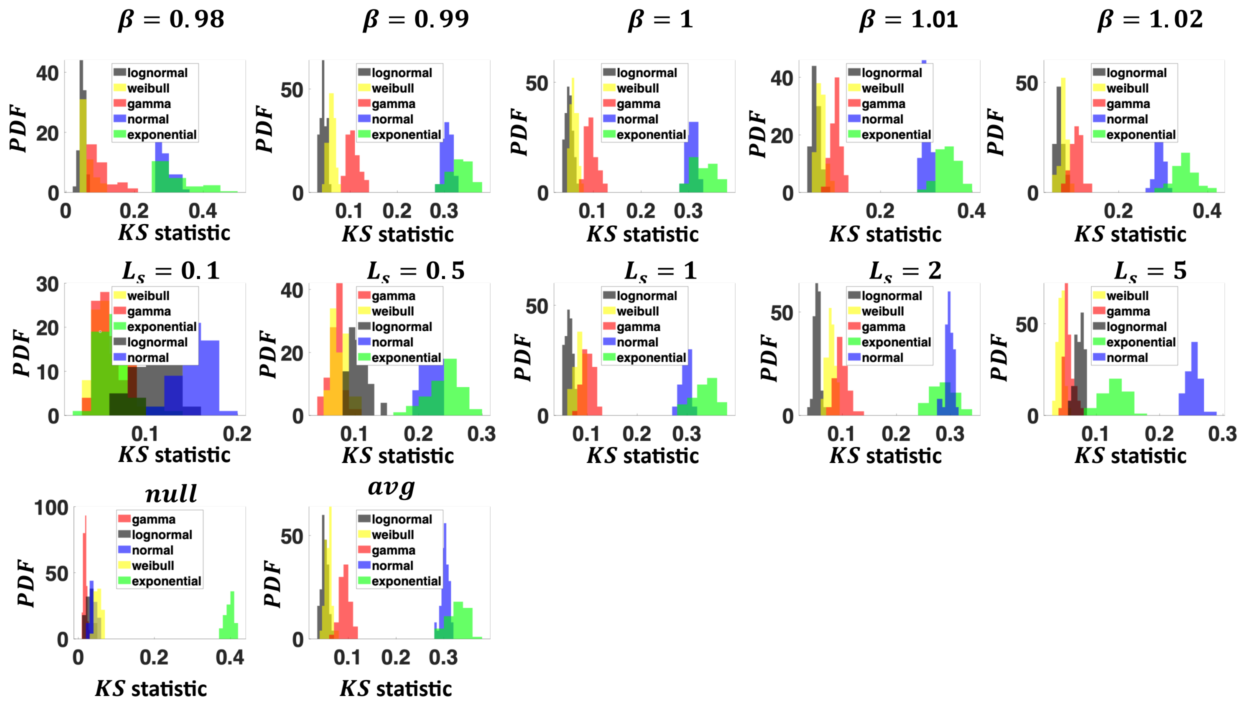


*Figure S5.* ***KS statistics of fitted candidate distributions.*** *Legends were ranked by ascending mean KS statics.*

It should be noted that connection weights in null networks were not best described as a log-normal distribution, despite the KS statistic is small (Fig.2 and Fig. S5). To better characterize the difference between model and null networks, distributions of normalized weights were shown in Fig. S6. Connection weights in null networks exhibited less variability (strong connections are rare) relative to in model networks.

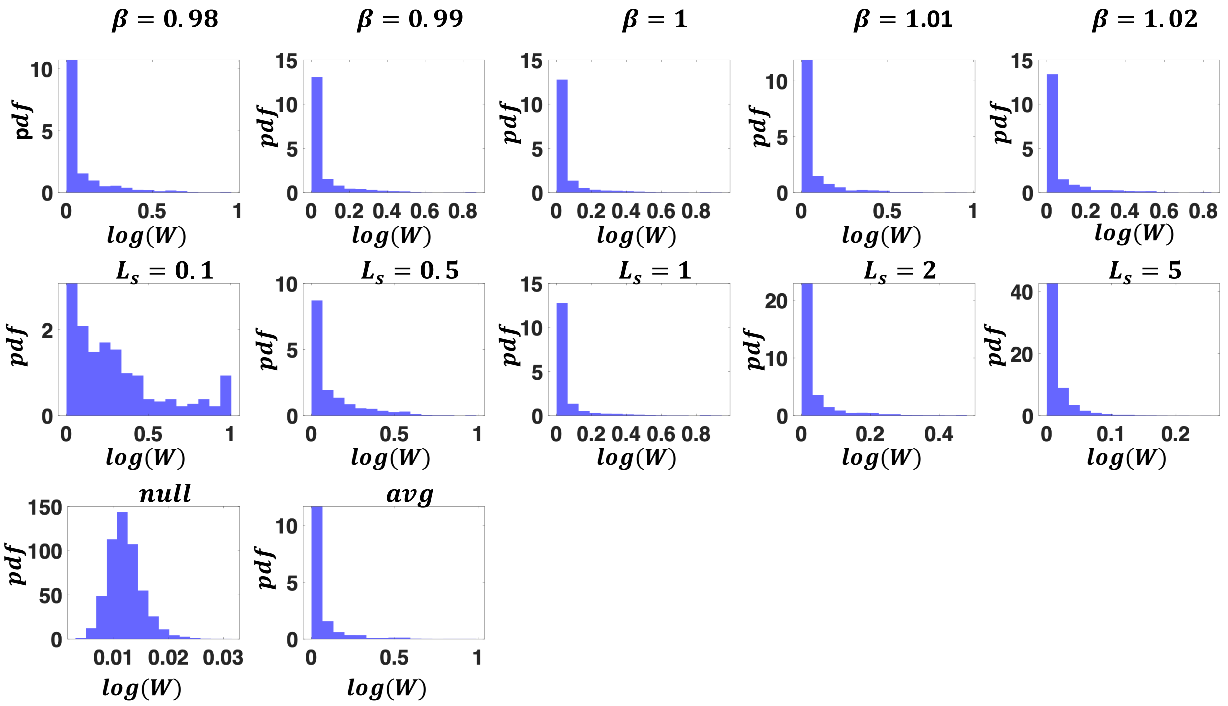


*Figure S6. **Probability density function of weight distributions (normalized by nodal strengths) in simulated networks.** Weights in null networks were small, denoting a lack of strong connections. In contrast, strong weights could be found in model networks.*

#### *p-value distributions of scale-free test*

In the main text, we suggested that the model was able to generate networks with scale-free degree distribution. The conclusion was drawn from the observations that more than 50% of networks generated by a parameter combination showed a $p > 0.1$ in the scale-free test. Fig. S7 showed a histogram of $p$-values in 1,000 generated networks, for each evaluated parameter combination. Null networks did not show a scale-free property. Scale-free was evident in $\beta =$ 0.98 and 1 ($L_s$ was fixed to 1), and all $L_s$ values considered ($\beta$ was fixed to 1). The group average parameters of HCP population were found to be able to generate scale-free networks, despite the p value marginally above the threshold of $p = 0.1$.

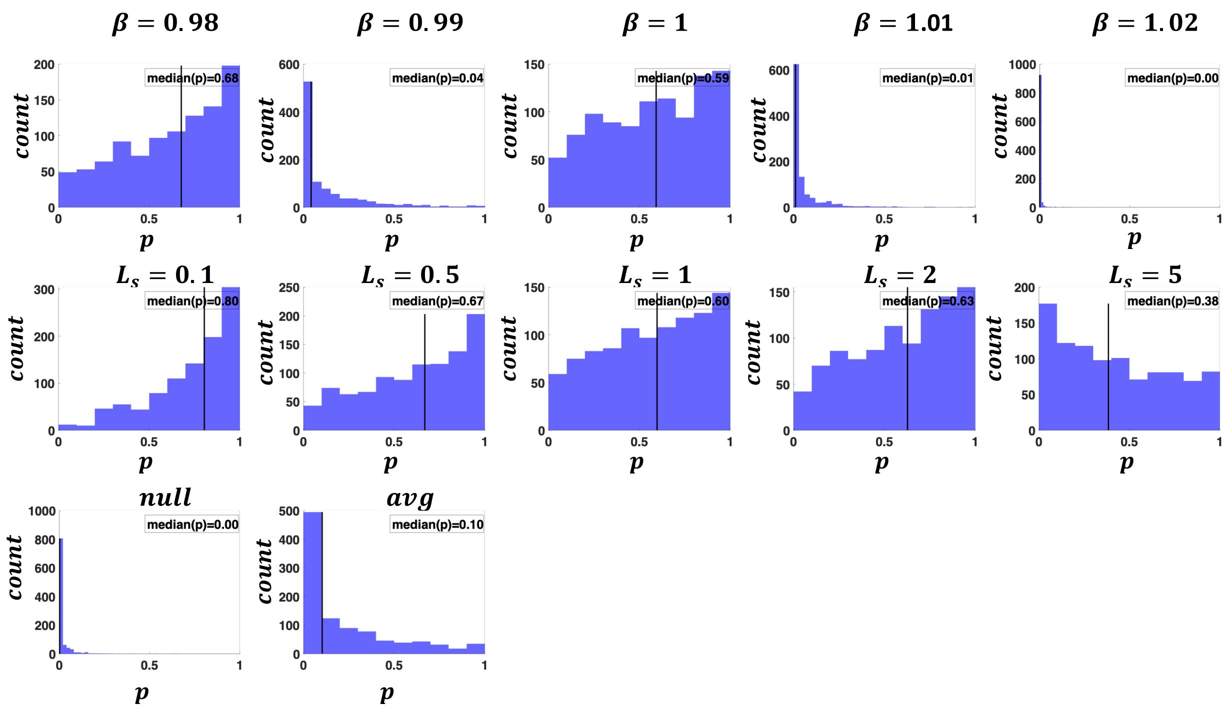


***Figure S7. p value distributions of scale-free tests.*** *Each histogram displayed p values of 1,000 networks.*

***Contour plot patterns of topological properties are insensitive to network density***

In Fig. 4, we evaluated the weighted topological properties of generated networks at a network density of 10%. We found that the patterns of contour plots are insensitive to network density and show the results for network density of 5% in Fig. S8.

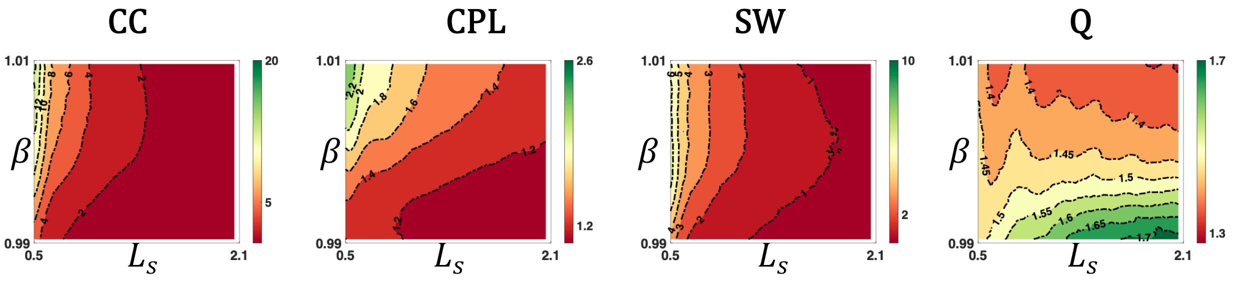


*Figure S8. Weighted topological properties (the same metrics as in Fig. 4a) evaluated at 5% network density.*

#### *Raw values of topological metrics*

In Fig. 4a, we showed the weighted topological properties of generated networks, normalized by weight and degree preserved null networks. Here in Fig. S9, we show the raw values of weighted topological properties of CC, CPL, and Q. SW is not included here because it is by definition normalized. It should be noted that modularity Q changed with parameters in a different way compared to Fig.4. This demonstrated that when $\beta$ is increasing (or $L_s$ is decreasing), modularity increases faster in null networks compared to model networks.

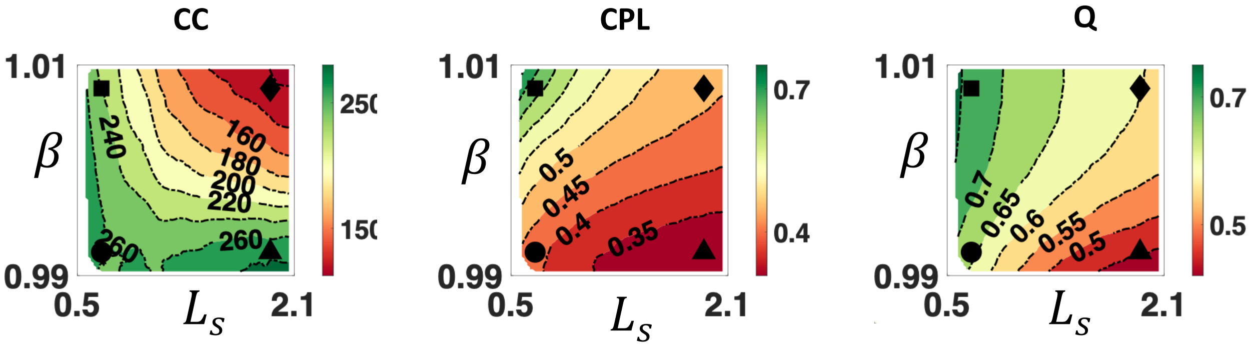


*Figure S9. Raw values of weighted topological properties (CC, CPL, and modularity Q).*

***Modified energy cost relative to the state-of-the-art models***

In this work, we developed a new parameter fitting cost function to fit the model parameters to individual connectomes. This is because the energy cost used in the current state-of-the-art models is not applicable to our model, due to the missing nodal correspondence between generated and empirical connectomes. However, three topological properties considered in the energy function, including the degree, clustering, and betweenness centrality, are still eligible for comparison. Thus, we modified the energy cost function by considering these three features only and compared the energy achieved by our model, compared to the state-of-the-art matching index model and pure geometric model.

As shown in Fig. S10, our model achieved a better fit compared to the geometric model, and a worse fit relative to the matching index model. This result is expected: The geometric model has the lowest model complexity (with only one model parameter). Although the matching index and our model both have 2 model parameters, our model relies on purely geometric information, where the matching index model used both geometric and topological information. In conclusion, geometry-dependent dynamic axon guidance generates networks better replicate empirical connectome than a simple distance rule, whereas this improvement is smaller than considering complex topological information.

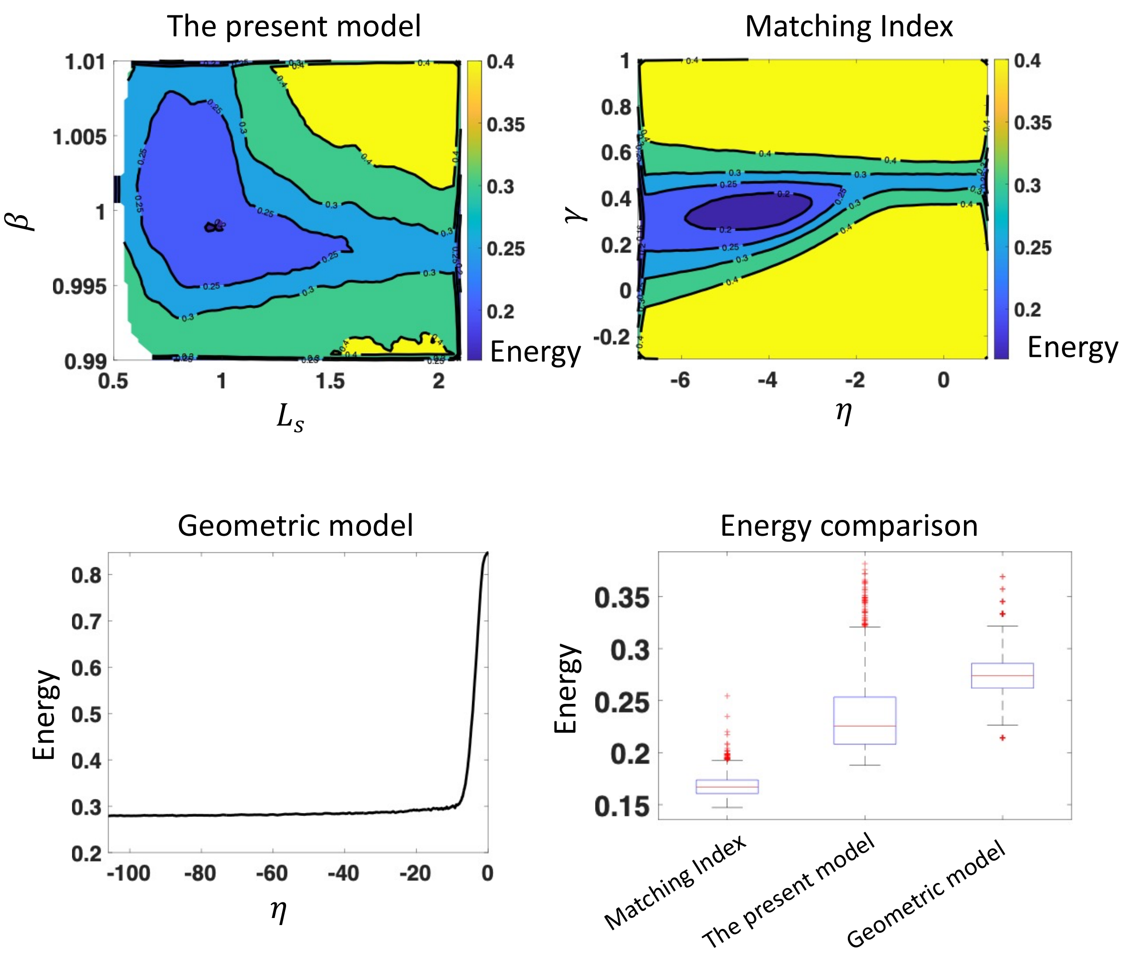


*Figure S10. Our model achieved a better energy compared to the state-of-the-art geometric model, and a worse energy compared to the state-of the-art matching index model. All energy functions consider degree, clustering, and betweenness centrality only.*

### *Parameter optimization: linear scaling and model fit*

As discussed above (the section explained the parameter specification of $N_a$), because axon sampling is probabilistic, the expectation of an edge weight should linearly scale with $N_a$. Thus, to optimize the model parameters, we linearly scaled the total connectivity of all networks (both model and empirical networks) to the same value.

A related concern is that this may disrupt the inter-individual variations in total connectivity. To test this, we randomly selected a sample of 100 participants from the HCP cohort. Model networks were linearly scaled to the total connectivity matching each individual before parameter optimization. We found that fitted parameters derived from the two approaches were comparable.

As per previous studies , we report the cost value of selected model networks in Fig. S11. Selected networks showed an RMSE of around 0.5 standard deviations on average, with comparable values among clustering coefficient, CPL, and modularity.

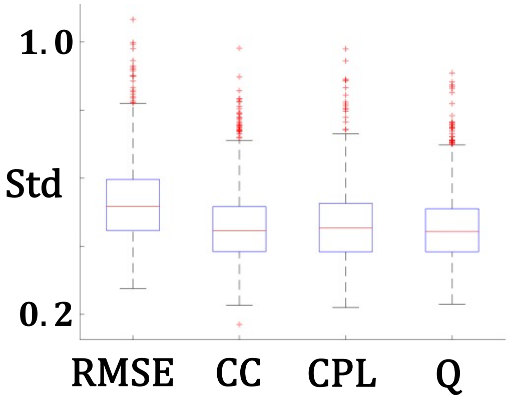


*Figure S11. RMSE, and cost in CC, CPL, and modularity Q, of networks selected to optimize parameters for the HCP cohort.*

***Associate model parameters to individual traits***

We optimized the model parameters for individual connectomes participating in the HCP dataset and associate them $(\beta, L_s)$ to age and sex. No significant association was found to age, probably because HCP is a healthy young adult dataset in which effects of development and aging are less evident. Males and females significant differ in $L_s$ (Fig. S12), suggesting model parameters can capture inter-individual variations in connectomes.

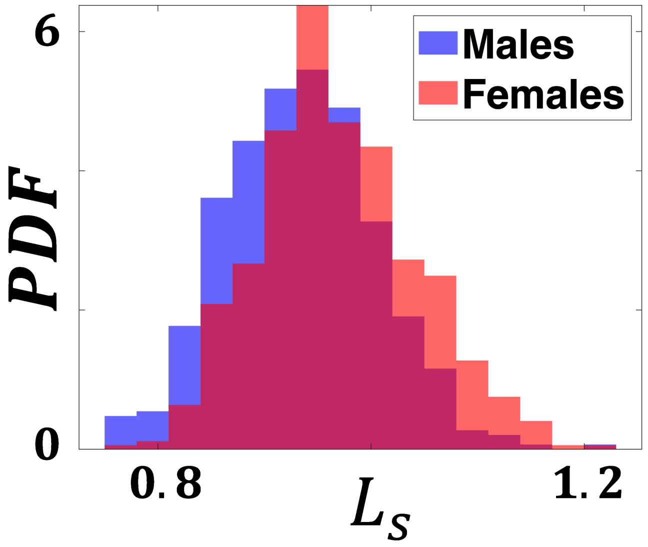


*Figure S12. Males and females significantly differ in $L_s$ ($p<0.05$, cohen's d = 0.49).*
